# Contrast thresholds reveal different visual masking functions in humans and praying mantises

**DOI:** 10.1101/135970

**Authors:** Ghaith Tarawneh, Vivek Nityananda, Ronny Rosner, Steven Errington, William Herbert, Sandra Arranz-Paraíso, Natalie Busby, Jimmy Tampin, Jenny Read, Ignacio Serrano-Pedraza

## Abstract

Recently, we showed a novel property of the Hassenstein-Reichardt detector: namely, that insect motion detection can be masked by “invisible” noise, i.e. visual noise presented at spatial frequencies to which the animals do not respond when presented as a signal. While this study compared the effect of noise on human and insect motion perception, it used different ways of quantifying masking in two species. This was because the human studies measured contrast thresholds, which were too time-consuming to acquire in the insect given the large number of stimulus parameters examined. Here, we run longer experiments in which we obtained contrast thresholds at just two signal and two noise frequencies. We examine the increase in threshold produced by noise at either the same frequency as the signal, or a different frequency. We do this in both humans and praying mantises (*Sphodromantis lineola*), enabling us to compare these species directly in the same paradigm. Our results confirm our earlier finding: whereas in humans, visual noise masks much more effectively when presented at the signal spatial frequency, in insects, noise is roughly equivalently effective whether presented at the same frequency or a lower frequency. In both species, visual noise presented at a higher spatial frequency is a less effective mask.

**Summary Statement:** We here show that despite having similar motion detection systems, insects and humans differ in the effect of low and high spatial frequency noise on their contrast thresholds.

## Introduction

Insect motion vision is thought to be mediated by arrays of retinotopic elementary motion detectors (EMDs) that cross-correlate the spatial input of a small region of the visual field with a similar, but temporally delayed, input of a neighbouring region. This cross-correlation model was first proposed by Hassenstein and Reichardt to describe the optomotor response of the beetle *Chlorophanus* (Hassenstein and Reichardt 1956) and has since demonstrated outstanding agreement with behavioural and neurophysiological observations across several forms of motion-elicited behaviour including tracking (Bahl et al. 2013), collision avoidance (Srinivasan et al. 1991) and landing (Borst and Bahde 1988). Experiments have elucidated the mechanisms underlying these responses (Borst 2014) and exposed the neural pathways that mediate detector computations (Riehle and Franceschini 1984) to a remarkable level of detail.

EMDs consist of two mirror-symmetrical subunits that compute motion in opposing directions (Fig). We recently showed (Tarawneh et al. 2017) that this opponency gives EMDs a surprising property: namely, motion detection can be impaired by noise which is “invisible” to the animal, in the sense that it does not elicit a response when presented as a signal. Our “signals” were drifting luminance gratings, i.e. a set of black and white vertical stripes moving horizontally across the screen. These signals, if detected, trigger the animal’s optomotor response, i.e. a body movement in the direction of the motion, which tends to keep the animal’s head aligned to the bars (Reichardt and Wenking 1969; Nityananda et al. 2015). The spatial frequency of the grating corresponds to the width of the bars: wide bars represent low spatial frequencies, narrow bars represent high. To add noise, we superimposed similar gratings, at either the same or different spatial frequency, which did not move smoothly but jumped around randomly. We found that low-frequency gratings did not elicit an optomotor response when presented on their own, moving smoothly, yet still disrupted the optomotor response to higher-frequency gratings when superimposed as noise. This disruption is referred to as “masking” of the signal by noise (Moore 2012; Anderson and Burr 1985).

We analysed the mathematics of the EMD to show why this effect occurs. Briefly, it works as follows. Insect ommatidia have a roughly Gaussian acceptance profile, so they effectively implement a low-pass spatial filter of the incoming light pattern. Thus, the first step in the EMD is a low-pass spatial filter (top row of Fig). Low spatial frequencies naturally elicit a response in these filters. However, the later opponency step (bottom row, “subtraction”, in Fig) cancels out this response for low enough spatial frequencies. This is why moving gratings at these frequencies do not elicit an optomotor response. However, when low frequencies are presented as noise superimposed on a signal grating of a higher frequency, this initial response can still act as noise, impairing the optomotor response elicited by gratings at higher frequencies.

Humans do not show this “invisible noise” effect. In our previous paper (Tarawneh et al. 2017), we compared our mantis data with previously-published human data and showed that there was a qualitative difference. In humans, noise is most effective when presented at the same frequency as the signal, and becomes less effective when presented either at higher *or* at lower frequencies than the signal (Anderson and Burr 1985). We were able to explain this too within the same mathematical framework: it arises because spatial filtering in humans is bandpass.

A limitation of our previous paper was that we did not compare exactly the same metrics of masking in humans and mantises. In humans, Anderson & Burr (Anderson and Burr 1985) had carried out extensive psychometric experiments measuring contrast thresholds at many different combinations of signal and noise. Their measure of masking was the ratio between the contrast threshold needed to judge the direction of a moving grating when it was presented alone, and the higher threshold needed when noise was added. They used data from 2 human observers, and contrast thresholds were obtained by the Method of Adjustment (i.e., observers adjusted the contrast of the moving grating by hand until its direction of drift was “just discernible”). In mantises, the Method of Adjustment is obviously not feasible. Previously (Nityananda et al. 2015), we have obtained mantis contrast thresholds from psychometric functions using the Method of Constant Stimuli (Lu and Dosher 2014). A human observer viewed mantises via a webcam and categorised each trial according to whether an optomotor response did or did not occur. The response rate increases with the contrast of the moving grating. The contrast threshold is then defined as the contrast at which a response occurs on half of the trials. The drawback of this method is that many trials are required. It was not feasible to run so many trials on each of many different combinations of signal and noise frequencies. Accordingly, in (Tarawneh et al. 2017) we used a different measure of masking in our 11 mantis observers. We kept the stimulus contrast fixed, and examined how the response rate varied for noise at different frequencies. We argued that these different metrics were equivalent, so should not affect our conclusions.

In this paper, we confirm this by running longer experiments for a subset of signal and noise combinations. Here, we use identical paradigms in both mantis and human observers as far as possible. As described above, stimuli are presented at a range of signal contrasts for a given noise condition, so as to obtain a complete psychometric function, which is then fitted to obtain the contrast threshold. In this independent data-set, we find the same qualitative difference in the effect of noise on human vs insect vision as with our earlier method. This confirms that the difference was not somehow an artefact of the masking metrics used in the earlier paper, and strengthens confidence in the result.

## Materials and Methods

### Human Experiment H1

#### Subjects

Data in Experiment H1 were collected from four subjects, all with experience in psychophysical experiments. Two were authors and two were naïve to the purposes of the study.

#### Visual Stimulus

The stimulus had “signal” and “noise” vertical sinusoidal gratings that moved either left or right in each trial. Signal gratings were of either 0.4 and 2 cpd frequencies and drifting at 8 Hz. Noise gratings were ideal band-pass filtered spatial noise with a bandwidth of 1 octave and a power spectral density of 0.02 (cpd)^-1^. The phase spectrum of the noise was updated randomly on every CRT frame (refresh rate was 60 Hz), making it temporally broadband up to the Nyquist temporal frequency of 30 Hz. The contrast levels of the signal and noise components were summed at each pixel (parameters were chosen to ensure that no clipping occurred). Each presentation lasted for 1 second. Still frames, space-time plots and spatiotemporal Fourier amplitude spectra of the masked condition stimuli used in Experiment H1 are shown in Fig 1.

#### Experimental Setup

Participants viewed stimuli on a 19” Eizo T765 CRT monitor from a distance of 100cm and indicated perceived direction of motion by keyboard presses. The monitor had a resolution of 1280×1024 pixels, 14-bit luminance levels and subtended a visual angle of 19.18×15.37 degrees at the viewing distance of participants. Its mean luminance was 57 cd/m^2^. Luminance was gamma corrected (gamma=2.31) using a Minolta LS-100 (Konica Minolta, Japan). A chin-rest (UHCOTech HeadSpot) was used to stabilize the subject’s head. Experiments were administered by a Matlab script using Psychophysics Toolbox Version 3. DataPixx Lite and ResponsePixx Handheld devices (VPixx Technologies Inc., Canada) were used to render stimuli and capture participant responses.

#### Experimental Procedure

Subjects indicated perceived direction of motion after each presentation and their contrast thresholds (per condition) were calculated using adaptive Bayesian staircases. Each staircase consisted of 50 trials and thresholds were averaged across three staircase repeats per condition for each subject.

### Mantis Experiments M1, M2

#### Insects

The insects used in experiments were 6 individuals of the species *Sphodromantis lineola*. Each insect was stored in a plastic box of dimensions 17×17×19 cm with a porous lid for ventilation and fed a live cricket twice per week. The boxes were kept at a temperature of 25° C and were cleaned and misted with water twice per week.

#### Visual Stimulus – Experiment M1

In Experiment M1, signal spatial frequencies (*f*_*S*_) were either 0.04 cpd or 0.2 cpd and noise was either (1) not present, (2) added with *f*_*n*_ = 0.04 cpd or (3) added with *f*_*n*_ = 0.2 cpd. Noise again had a spatial bandwidth of 1 octave. For each of the 6 combinations of grating frequency and noise setting, trials were run with grating Michelson contrast levels of [2^-6^, 2^-5^ … 2^-1^] to calculate contrast detection thresholds. There were thus 36 different conditions in total. A total of 6 mantises each ran 10 repeats of each condition (360 trials).

#### Visual Stimulus – Experiment M2

Ideally, we wished our grating to contain a single spatial frequency when presented on the mantis retina. A grating rendered on a flat screen whose spatial periods are constant in pixels is non-uniform in visual degrees (Anderson and Burr 1987): a given number of pixels at the edge of the screen projects to a smaller angle than the same number directly in front of the viewer. This distortion is generally neglected in human psychophysics but is potentially important at the small viewing distance (7 cm) used in our experiments. To correct for this, we applied a non-linear horizontal transformation in Experiment M2 so that grating periods subtend the same visual angle irrespective of their position on the screen, using the technique described in Nityananda et al. 2017. This was achieved by calculating the visual degree corresponding to each screen pixel using the function:

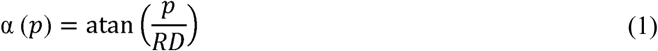

where *p* is the horizontal pixel position relative to the centre of the screen, α (*p*) is its visual angle, R is the horizontal screen resolution in pixels/cm and *D* is the viewing distance. To an observer standing more than *D* cm away from the screen, a grating rendered with this transformation looked more compressed at the centre of the screen compared to the periphery. At *D* cm away from the screen, however, grating periods in all viewing directions subtended the same visual angle and the stimulus thus appeared uniform (in degrees) as if rendered on a cylindrical drum. This correction only works perfectly if the mantis head is in exactly the intended position at the start of each trial, and is most critical at the edges of the screen. As an additional precaution against spatial distortion or any stimulus artefacts caused by oblique viewing we restricted all gratings to the central 85° of the visual field by multiplying the stimulus luminance levels *L*(*x*, *y*, *t*) with the following Butterworth window:

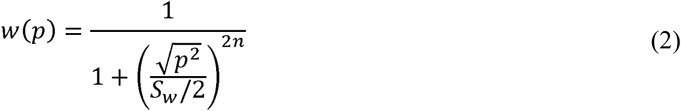

where *x* is the horizontal pixel position relative the middle of the screen, *S_w_* is the window size (distance between the 0.5 gain points) in pixels, chosen as 512 pixels in our experiment (subtending a visual angle of 85° at the viewing distance of the mantis) and *n* is the window order (chosen as 10). This restriction minimised any spread in spatial frequency at the mantis retina due to imperfections in our correction formula described by Equation (1).

With the above manipulations the presented stimulus (as a function of pixel horizontal position *p* and frame number *j*) was:

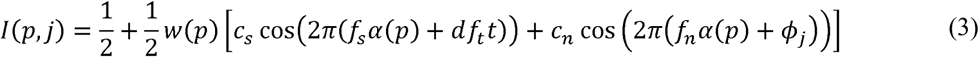

where *I* is pixel luminance, in normalised units where 0 is the minimum and 1 the maximum luminance available on the screen (0.161 and 103 cd/m^2^ respectively), *c*_*s*_ is signal contrast (varied across trials), *c*_*n*_ is noise contrast (fixed at 0.2), *f*_*S*_ and *f*_*n*_ are the spatial frequency of signal and noise respectively (both varied across trials), *f*_*t*_ is signal temporal frequency (fixed at 8 Hz), *d* indicates motion direction (either 1 or -1 on each trial), *φ*_*j*_ is picked randomly from a uniform distribution between 0 and 1 on every frame of the trial, *t* is time in seconds (given by *t* = *j*/85), and *α*(*p*) is the pixel visual angle according to Equation (1).

Still frames, space-time plots and spatiotemporal Fourier amplitude spectra of the masked condition stimuli used in Experiment M2 are shown in Fig.

#### Experimental Setup

The setup consisted of a CRT monitor (HP P1130) and a 5×5 cm Perspex base onto which mantises were placed hanging upside down facing the [horizontal and vertical] middle point of the screen at a distance of 7 cm. The Perspex base was held in place by a clamp attached to a retort stand and a web camera (Kinobo USB B3 HD Webcam) was placed underneath providing a view of the mantis but not the screen. The monitor, Perspex base and camera were all placed inside a wooden enclosure to isolate the mantis from distractions and maintain consistent dark ambient lighting during experiments.

The screen had physical dimensions of 40.4×30.2 cm and pixel dimensions of 1600×1200 pixels. At the viewing distance of the mantis the horizontal extent of the monitor subtended a visual angle of 142°. The mean luminance of the monitor was 13.2 cd/m^2^ and its refresh rate was 85 Hz.

The monitor was connected to a Dell OptiPlex 9010 (Dell, US) computer with an Nvidia Quadro K600 graphics card and running Microsoft Windows 7. All experiments were administered by a Matlab 2012b (Mathworks, Inc., Massachusetts, US) script which was initiated at the beginning of each experiment and subsequently controlled the presentation of stimuli and the storage of keyed-in observer responses. The web camera was connected and viewed by the observer on another computer to reduce the processing load on the rendering computer’s graphics card and minimize the chance of frame drops. Stimuli were rendered using Psychophysics Toolbox Version 3 (PTB-3) (Brainard 1997; Pelli and Brainard 1997; Kleiner et al. 2007).

#### Experimental Procedure

Each experiment consisted of a number of trials in which an individual mantis was presented with moving gratings of varying parameters. An experimenter observed the mantis through the camera underneath and blindly coded the direction of the elicited optomotor response (if any). The response code for each trial was either “moved left”, “moved right” or “did not move”. There were equal repeats of left-moving and right-moving gratings of each condition in all experiments. Trials were randomly interleaved by the computer.

In between trials a special “alignment stimulus” was presented and used to steer the mantis back to its initial body and head posture as closely as possible. The alignment stimulus consisted of a chequer-like pattern which could be moved in either horizontal direction by keyboard shortcuts and served to re-align the mantis by triggering the optomotor response.

#### Calculating Contrast Detection Thresholds

After conducting Experiments M1 and M2 we calculated motion probability *P* (for each individual and stimulus condition) as the proportion of trials in which the mantis moved in the same direction as the signal grating. As in (Nityananda et al. 2015), the number of trials on which the mantis was coded as moving in the opposite direction was negligible. We then fitted the individuals’ responses using the psychometric function:

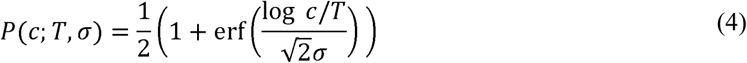

where *c* is the contrast of the signal grating, *T* is the contrast detection threshold (the contrast corresponding to *P* = 0.5) and *σ* represents the function’s steepness. The parameters *c*_*th*_ and *σ* were calculated using maximum likelihood estimation in Matlab assuming the mantises’ responses had a simple binomial distribution. Therefore:

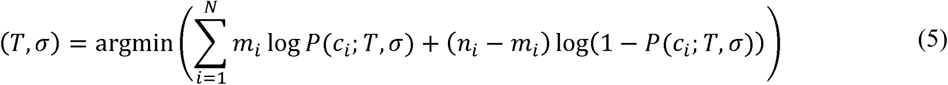

where the subscript *i* indicates different contrast levels, *n*_*i*_ is the total number of trials done for contrast *c*_*i*_, *m*_*i*_ is the number of trials in which the mantis moved in the grating direction, *p* is motion probability given by Equation (4) and *N* is the number of contrast levels. Detection thresholds were fitted to each insect’s individual data, and detection thresholds were then averaged across the population.

#### Data Collection

In Experiment M1, six animals ran two blocks of trials each. Each block had 360 randomly-interleaved trials consisting of 60 repeats of each grating condition and mantis responses were coded by WH.

In Experiment M2, six animals ran a single block of trials each. Each block had 360 randomly-interleaved trials consisting of 60 repeats of each grating condition and mantis responses were coded by SE.

## Results

### Experiment H1

We measured the contrast detection thresholds of 4 human observers for direction discrimination in moving gratings under different masking conditions. The signal was a vertical sinusoidal grating (of temporal frequency 8 Hz, spatial frequency *f*_*S*_ and variable contrast) drifting to either left or right in each presentation. We used two different signal frequencies: *f*_*S*_ = 0.4 cpd and *f*_*S*_ = 2 cpd. Pilot work indicated that these were on either side of peak sensitivity and that thresholds were the same for both. Noise was added in a subset of trials; it had a spatial bandwidth of 1 octave around either 0.4 or 2 cpd and was temporally broadband.

We will henceforth refer to the various conditions of our masking experiments using the notation S+N where S indicates signal frequency and is either H for the high frequency (2 cpd) or L for the low frequency (0.04 cpd) and N similarly indicates noise frequency. The condition H+L therefore refers to the grating with *f*_*S*_ = 2 cpd and *f*_*n*_ = 0.4 cpd. Grating conditions with no noise are simply referred to as H and L. Still frames, space-time plots and the spatiotemporal Fourier amplitude spectra of the masked conditions (L+L, L+H, H+L, H+H) are shown in Fig.

Human contrast thresholds are shown in Fig 2. The thresholds for signal alone (H and L) do not differ significantly. Adding noise at either frequency caused a significant increase in threshold for both signal frequencies: there were significant differences in thresholds for L+L and L (paired t(3) = 13.0, p < 0.001), for L+H and L (paired t(3) = 13.8, p < 0.001), for H+L and H (paired t(3) = 14.7, p < 0.001) and for H+H and H (paired t(3) = 26.0, p < 0.001). However, for both the 0.4 and 2 cpd signal frequencies, the threshold was higher when noise and signal frequencies were the same compared to when they were different: there were significant differences in thresholds for L+L and L+H (paired t(3) = 4.5, p = 0.021) and for H+H and H+L (paired t(3) = 8.4, p = 0.003). These results are consistent with previous studies in human literature which have shown that maximal masking occurs when noise is of equal or near frequency to that of the signal (Anderson and Burr 1985, 1989).

### Experiment M1

We also ran essentially the same experiment in insects. Mantises were placed in front of a computer screen and viewed full screen gratings drifting horizontally to either left or right in each trial. The stimuli were the same as described for humans above, except that the spatial frequencies were lowered in order to account for the poorer spatial acuity of insect vision. The high (H) and low (L) spatial frequencies were set to 0.2 and 0.04 cpd respectively (the mantis optomotor response is approximately equally sensitive at those frequencies (Nityananda et al, 2015)). An experimenter observed the mantis through a camera and coded the direction of the elicited optomotor response (if any) blind to the stimulus. Detection rates were later calculated per condition and individual as the proportion of trials in which the mantis moved in the same direction as the grating.

We measured the contrast detection thresholds for each of the conditions L+L, L+H, H+L, H+H as well as the non-masked grating conditions H and L. Fig 3 shows the response rates and fitted psychometric functions of one individual for illustration. Clearly, adding noise tends to move the psychometric function to the right. That is, in the presence of noise, the signal grating has to have higher contrast before it will reliably elicit an optomotor response from the insect. To quantify this we compared the contrast detection thresholds, averaged across 6 mantises, for the 6 different conditions (Fig 4). In the absence of noise, thresholds do not differ significantly between the low and high spatial frequency of the signal gratings (paired t(5) = 0.7, p = 0.494 comparing H and L) as expected since these two frequencies were chosen to drive the optomotor response equally. Adding low-frequency noise significantly increases thresholds for both signal frequencies: there were significant threshold differences for L+L and L (paired t(5) = 7.9, p < 0.001) and for H+L and H (paired t(5) = 4.8, p = 0.005). We again see no difference in thresholds depending on the signal frequency (paired t(5) = 2.0, p = 0.096 comparing H+L and L+L). However, when we add high-frequency noise, we see a very large difference between the two signal frequencies. High-frequency noise again significantly increases thresholds (paired t(5) = 4.0, p = 0.01 comparing L+H and L, paired t(5) = 7.6, p < 0.001 comparing H+H and H). The high-frequency signal is affected as badly by high-frequency noise as by low-frequency noise (paired t(5) = 0.1, p = 0.894 comparing H+H and H+L). However, the low-frequency signal is far less affected by high-frequency noise (paired t(5) = -4.2, p = 0.009 comparing thresholds for L+H and L+L). Note that this is not because high-frequency noise has an intrinsically small effect. The high-frequency noise has a very substantial effect on the high-frequency signal, just not on the low-frequency signal.

Experiments H1 and M1 demonstrate the presence of interactions between signal and noise frequencies in both humans and mantises. The responses of the two species, however, were qualitatively different. In humans, noise had a greater effect when presented at the signal frequency and a lesser effect at the other frequency. Mantises on the other hand were affected to the same degree by either noise frequency at the 0.2 cpd signal frequency, and more strongly by the noise frequency 0.04 cpd when signal frequency was also 0.04 cpd. In other words, mantises were affected most when noise frequency was equal or lower than signal frequency (across the frequencies 0.04 and 0.2 cpd). This indicates a qualitative difference between the two species.

### Experiment M2

The stimuli and experimental procedures were as similar as possible for both humans and mantises, with spatial frequencies chosen appropriate to each species’ contrast sensitivity function. However, one difference was that mantises were observing the screen from a much shorter viewing distance (7 cm as opposed to 100 cm for human subjects). When viewing a flat screen from a short distance the stimulus appears spatially distorted; uniform gratings subtend smaller visual angles at the periphery and may therefore consist of several spatial frequencies (in cpd). Thus, for mantises, the signal gratings effectively varied in spatial frequency across the stimulus, whereas for humans they were much more nearly constant. To test whether this distortion could have influenced our findings from Experiment M1, we repeated the same experiment using a modified stimulus. Previous studies have shown that the optomotor response of the mantis is driven predominantly by the central visual field (Nityananda et al 2017). The new stimulus was therefore different in three ways: (1) it was limited to the central 85 degrees of the visual field, (2) it was corrected for spatial distortion by introducing a non-linear horizontal transformation and (3) noise was restricted to a single spatial frequency.

Fig 5 shows the mean of the contrast detection thresholds, averaged across the 6 insects, for the six different conditions. Sensitivity was now much lower, particularly for the high frequency, presumably reflecting the alterations to the stimulus. Despite these differences, we found the same qualitative trend observed in Experiment M1. Masking was strongest when noise frequency was equal to or lower than signal frequency (across 0.04 and 0.2 cpd). The addition of noise caused a significant increase in thresholds across all conditions: L+L and L (paired t(5) = 12.5, p < 0.001), L+H and L (paired t(5) = 8.7, p < 0.001), H+L and H (paired t(5) = 3.8, p = 0.013) and H+H and H (paired t(5) = 2.7, p = 0.043). For the 0.04 cpd signal frequency grating, noise at the same frequency caused a significantly larger increase compared to noise at the higher frequency (paired t(5) = 6.4, p < 0.001 comparing L+L and L+H). There was no significant difference, however, between adding noise at either frequency in case of the 0.2 cpd signal frequency (paired t(5) = 1.1, p = 0.324 comparing H+L and H+H). That is, noise is equally effective whether added at the signal frequency or at a lower frequency, but less effective when added at a higher frequency. The agreement between our findings in Experiments M1 and M2 suggest that the difference in stimulus viewing distances, and the resultant distortion, does not explain the qualitative differences in mantis and human responses.

## Discussion

In a previous paper, we documented a striking difference between insect and human motion perception (Tarawneh et al. 2017). This difference relates to the robustness of perception under visual noise. In both species, some spatial frequencies are more effective “masks” than others. In human vision, the effectiveness of a given spatial frequency as a mask depends on two things: (i) how visible that spatial frequency is to the organism, and (ii) how close that spatial frequency is to the signal. Noise at frequencies that are less visible, or that are further from the signal frequency, is less effective at masking the signal. In insects, this apparently obvious result does not hold. We found previously that while noise higher than the signal frequency does indeed lose effectiveness as a mask, noise lower than the signal masks essentially independently of the distance between noise and signal – even when the noise is presented at such low frequencies that a signal there would elicit no response.

Because this finding is so counter-intuitive, it was important to validate it with a different paradigm, in which we directly compare humans and insects. That is what we have done in the present paper. Here, we selected two spatial frequencies, one high and one low, on either side of the organism’s peak sensitivity. These were chosen to be equally visible, or more precisely, to have equal contrast thresholds on a motion direction discrimination task. We then examined the effect of adding noise either at the same frequency, or the other frequency. We measured thresholds using the psychometric function obtained with the Method of Constant Stimuli, with the contrast of the signal grating as the varying parameter (Fig 3).

In humans, noise had a significantly stronger masking effect when at the same frequency as the signal; noise at a higher or lower frequency had less effect (Fig 2). In mantises, noise was equally effective whether added at the signal frequency or at a lower frequency, but less effective when added at a higher frequency (Fig 5). This agrees with our previous experiment using response rates at a fixed contrast. The effect is predicted by the current model of insect motion perception, combined with the low-pass nature of insect spatial filtering (Tarawneh et al. 2017). The experiments presented here confirm that the phenomenon affects contrast threshold, not just the response rate at one selected contrast, and strengthen our previous conclusions.

## Acknowledgements

Thanks to David Peterzell, Candy Rowe and Claire Rind for helpful discussions. Thanks to Adam Simmons for excellent insect care and husbandry.

### Competing Interests

The authors declare they have no competing interests

### Funding

VN, GT and RR were supported by Leverhulme Trust Research Leadership Award RL-2012- 019 to JR. ISP was supported by grant PSI2014-51960-P from Ministerio de Economía y Competitividad (Spain).

### Data Availability

Data will be made publicly available on at www.jennyreadresearch.com.

### Author Contributions

GT/ISP/VN: designed the experiments

GT/ISP: programmed visual stimuli and experiment scripts

GT: analysed the results and wrote the paper

SAP: carried out the human experiment (H1)

SE/WH: carried out mantis experiments (M1, M2)

VN/RR: contributed to discussions and helped write paper

NB/JT: conducted pilot experiments

JR/ISP: supervised the project and helped write paper

## Figures

**Fig 1:**
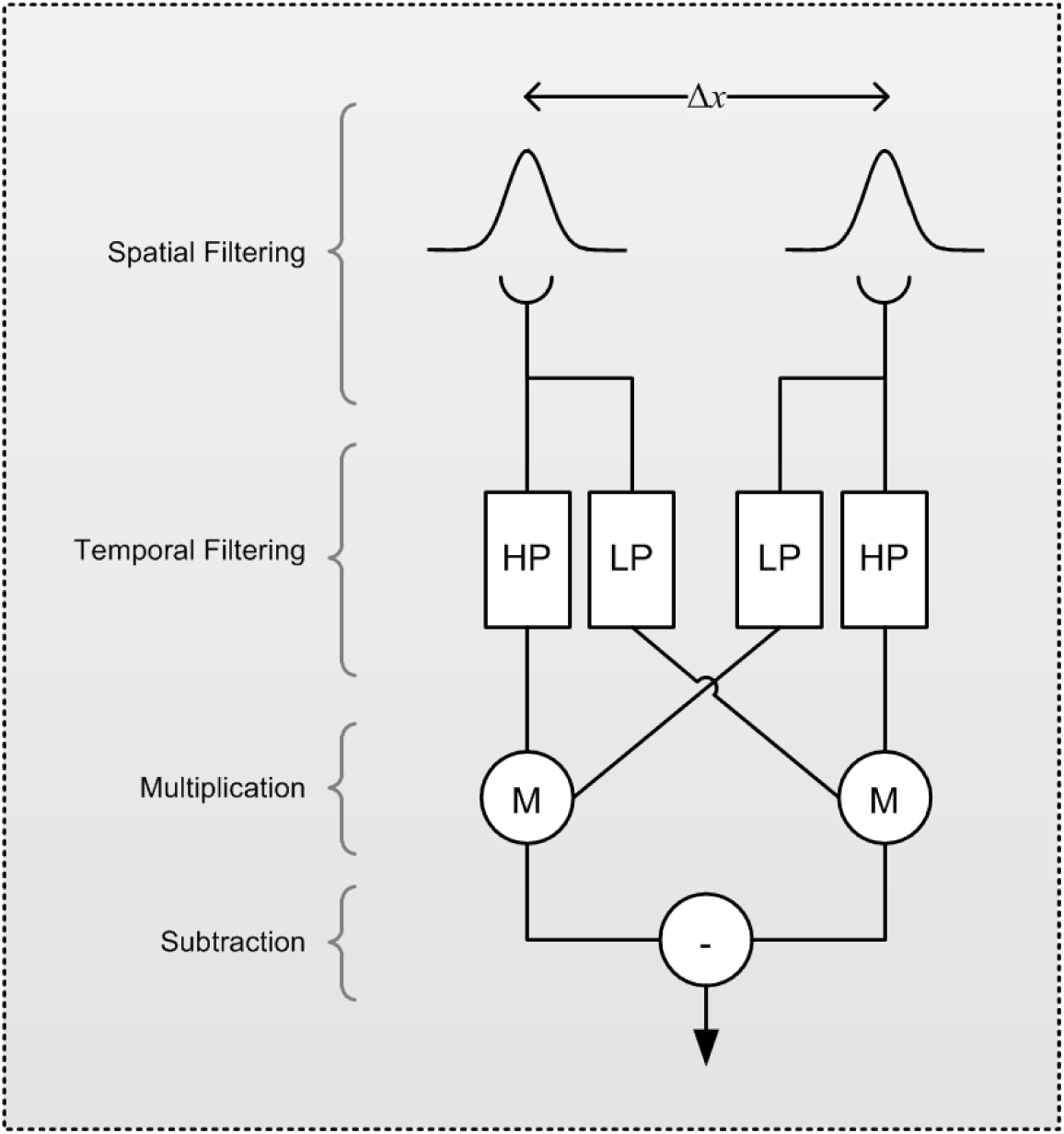
The Elementary Motion Detector (EMD). The spatial input from two identical Gaussian filters separated by Δ*x* is passed through high and low pass temporal filters (HP and LP respectively). The LP output in each subunit is cross-correlated with the HP output from the other subunit using a multiplication stage (M) and the two products are then subtracted to produce a direction-sensitive measure of motion.

**Fig 2:**
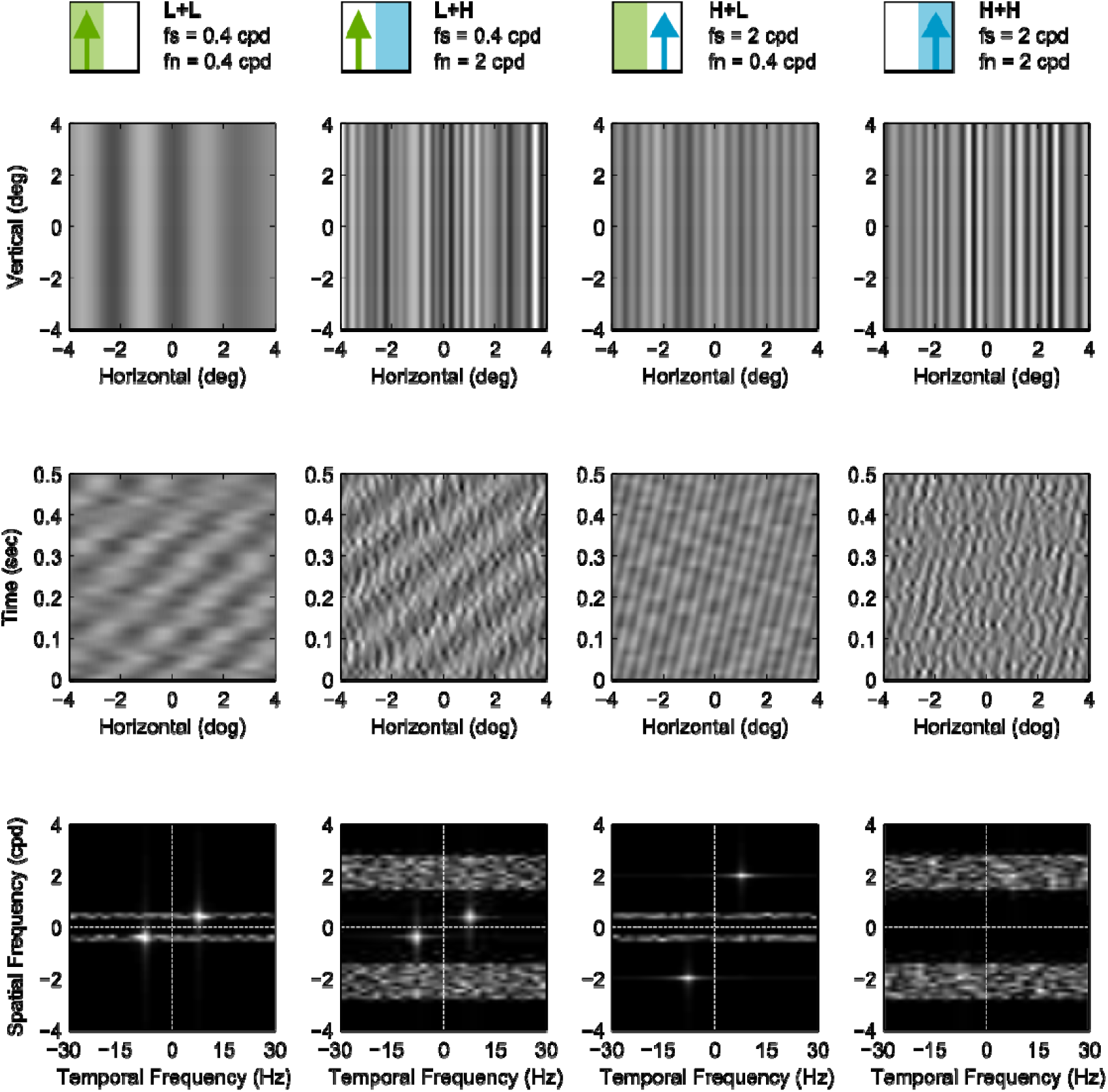
Masked grating stimulus conditions used in Experiment H1. Each column of panels represents one stimulus condition. The cartoons at the top represent the conditions graphically and are used in subsequent figures for easy reference (signal is the upwards pointing arrow and noise is the coloured rectangle). Top row of panels shows still frames of each condition while middle and bottom panel rows show corresponding space-time plots and Fourier spatio-temporal amplitude spectra respectively. In these plots the signal contrast was set to 0.2.

**Fig 3:**
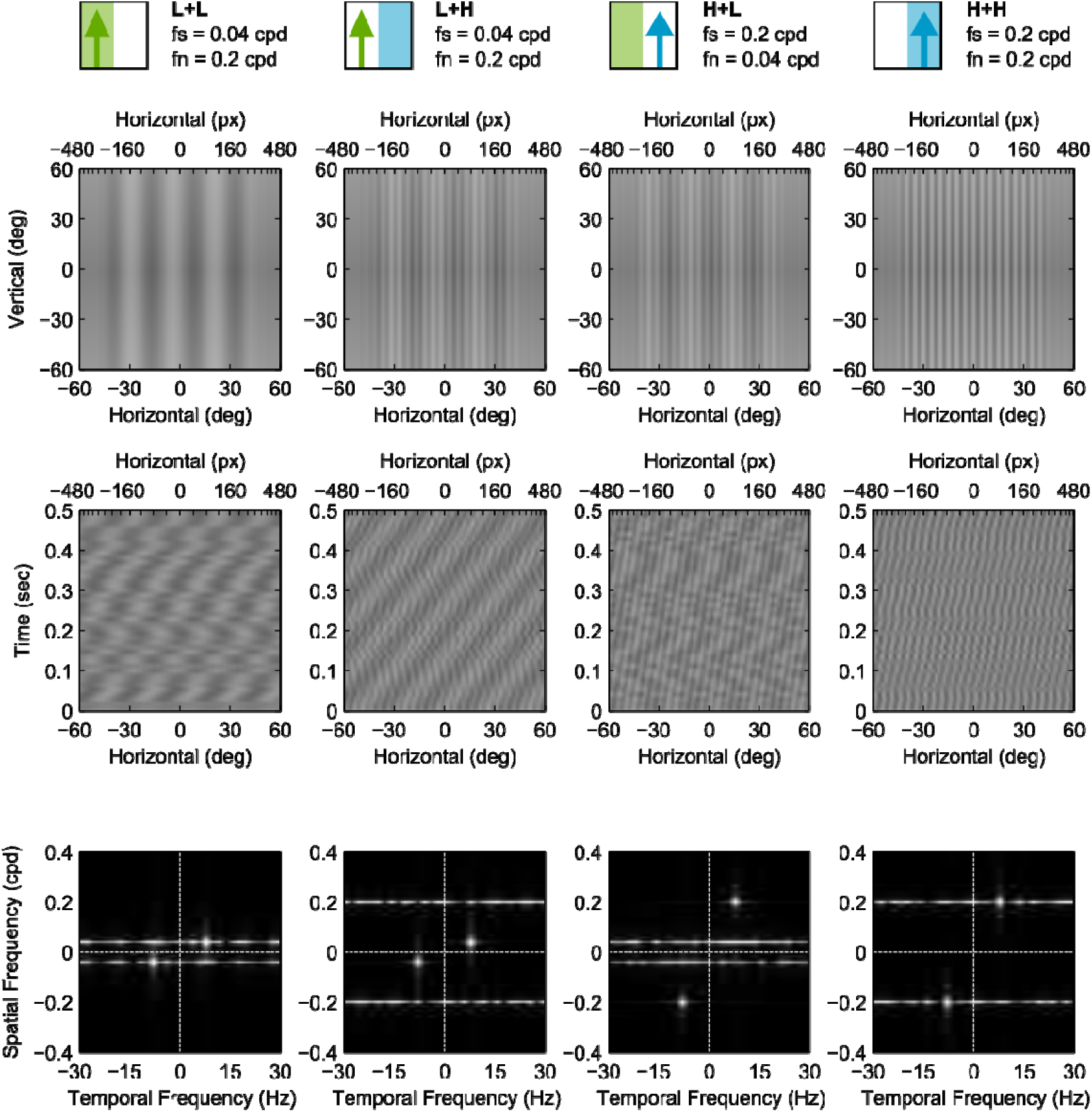
Masked grating stimulus conditions used in Experiment M2. Each column of panels represents one stimulus condition. Top row of panels shows still frames of each condition while middle and bottom panel rows show corresponding space-time plots and Fourier spatio-temporal amplitude spectra respectively. In these plots the signal contrast was set to 0.1. These stimuli conditions are similar to their correspondents in Experiment H1 (Fig) but were modified in three ways: (1) they were limited to the central 85 degrees of the visual field, (2) they were corrected for spatial distortion by introducing a non-linear horizontal transformation and (3) their noise was restricted to a single spatial frequency.

**Fig 2:**
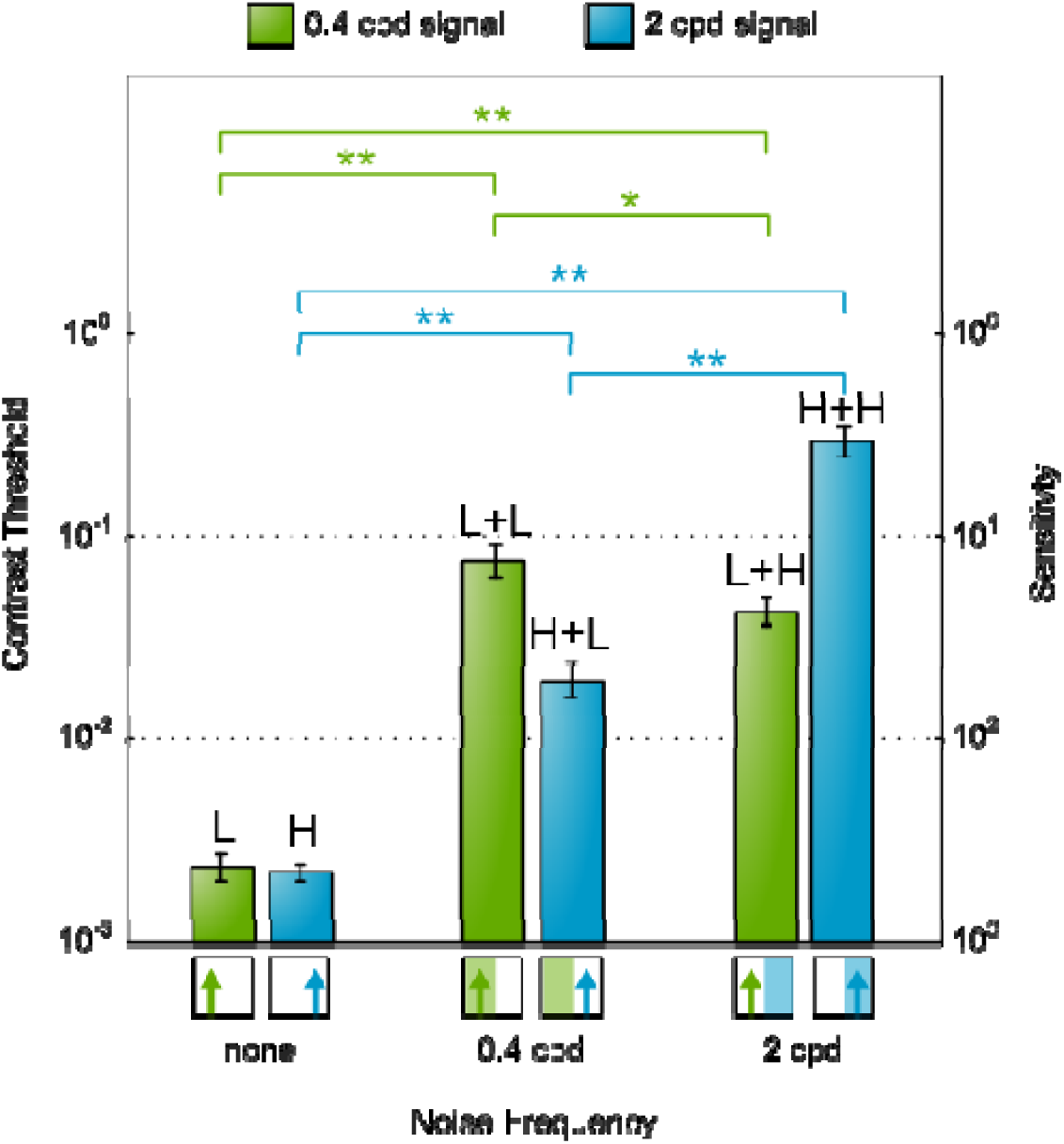
Human motion detection contrast thresholds for different combinations of signal and noise frequencies (measured in Experiment H1). Bars show mean contrast detection thresholds for 4 subjects and error bars show ± standard error of the mean. Horizontal brackets indicate threshold pairs that differ significantly (paired t-test, * p ≤ 0.05 and ** p ≤ 0.01). Results show that each of the two signals frequencies 0.4 (blue) and 2 cpd (green) was masked significantly higher by same-frequency noise compared to different-frequency noise. Letters above bars indicate signal and noise conditions in the format Signal + Noise (S + N). L indicate low spatial frequency, H indicates high spatial frequency. So for example L + H indicates low frequency signal and high frequency noise.

**Fig 3:**
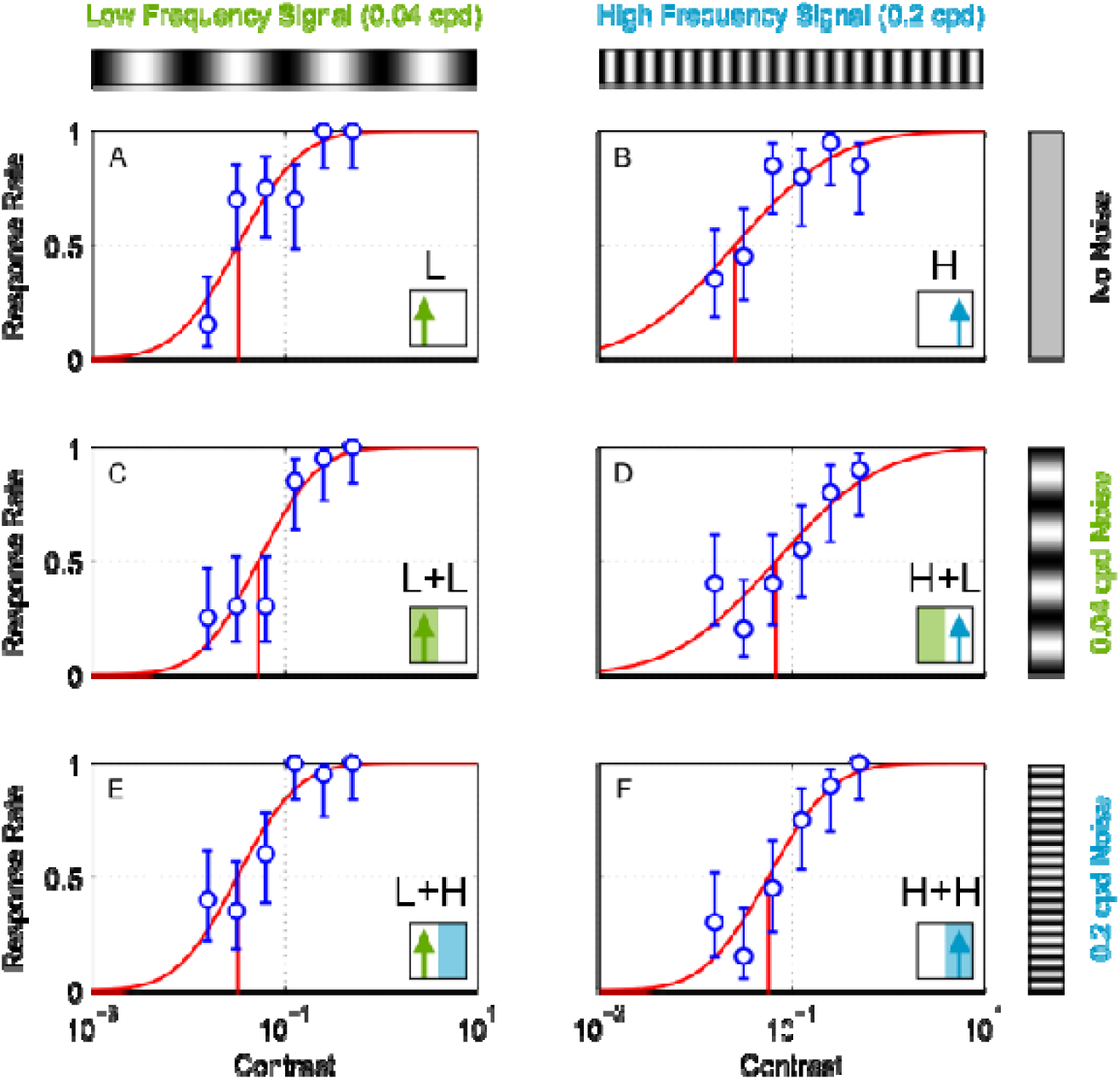
Responses, fitted psychometric curves and detection thresholds of a single mantis (measured in Experiment M1). Circles show response rates (i.e. proportion of trials on which the mantis was coded as moving in the same direction as the signal grating) as a function of signal grating contrast. Error-bars are 95% confidence intervals calculated from simple binomial statistics. Red curves show fitted psychometric function (Equation (4)); red vertical lines mark contrast threshold. (ACE): low-frequency signal (i.e. 0.04 cpd); (BDF): high-frequency signal (i.e. 0.2 cpd). Insets at the bottom right corner of each panel indicate signal and noise frequencies as in Fig 2. (AB): No noise: stimulus is a pure drifting luminance grating. (CD): Low-frequency noise, i.e. superimposed on the drifting signal grating is a grating of 0.04 cpd whose phase is updated randomly on every frame. (EF): High-frequency noise. The data plotted in this figure are all from a single individual (mantis F11) and were measured in Experiment M1. Letters to the right of the curves indicate signal and noise conditions in the format Signal + Noise (S + N). L indicate low spatial frequency, H indicates high spatial frequency. So for example L + H indicates low frequency signal and high frequency noise.

**Fig 4:**
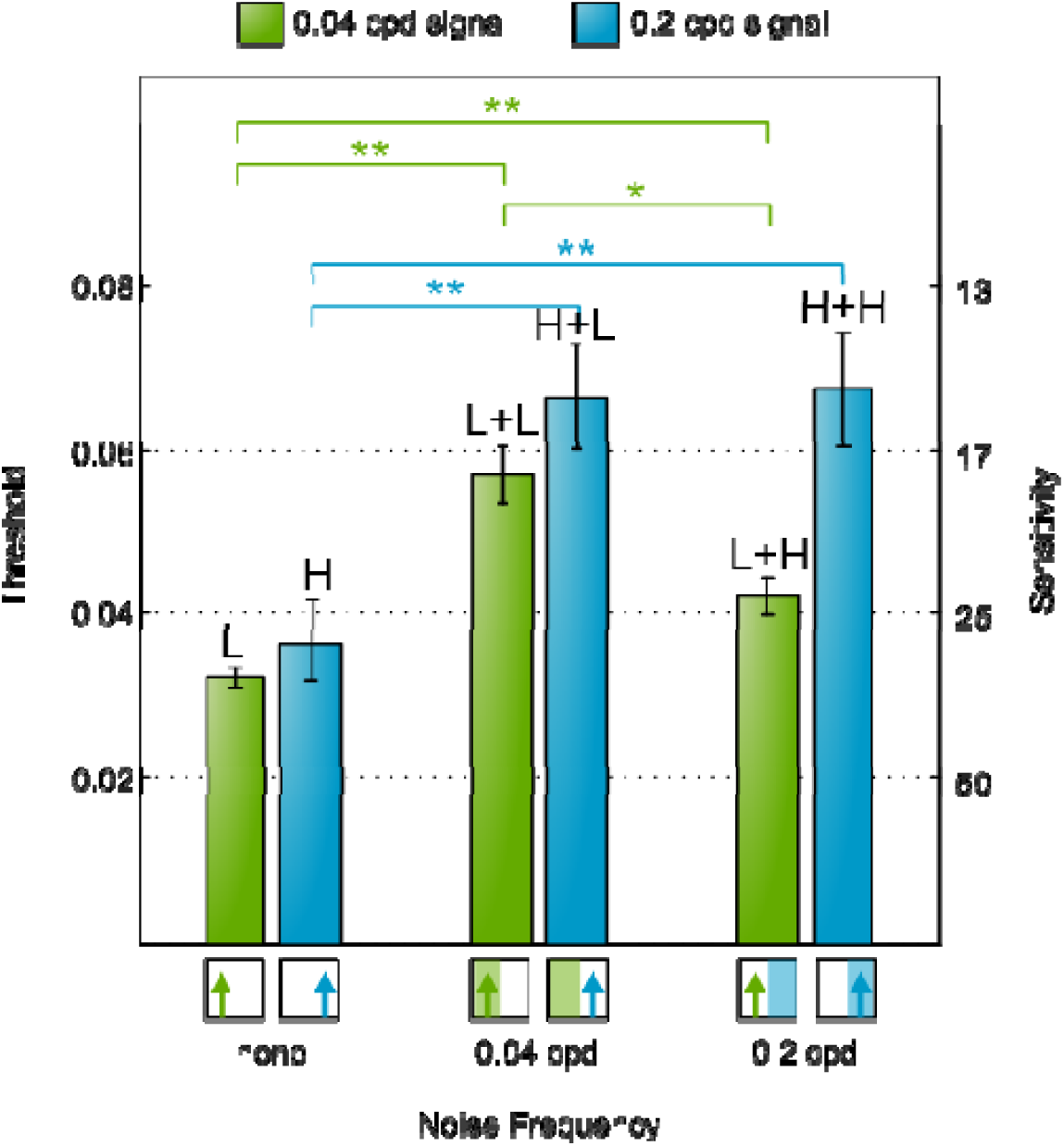
Mantis motion detection contrast thresholds for different combinations of signal and noise frequencies (measured in Experiment M1). Bars show mean contrast detection thresholds for 6 insects and error bars show ± standard error of the mean. Horizontal brackets indicate threshold pairs that differ significantly (paired t-test, * p ≤ 0.05 and ** p ≤ 0.01). Diagrams below bars indicate stimuli conditions as in Fig 2. Results show that the 0.2 cpd signal was masked to similar degrees by noise at either frequency while the 0.04 cpd signal was masked more strongly by the 0.04 cpd noise. Letters above bars indicate signal and noise conditions in the format Signal + Noise (S + N). L indicate low spatial frequency, H indicates high spatial frequency. So for example L + H indicates low frequency signal and high frequency noise.

**Fig 5:**
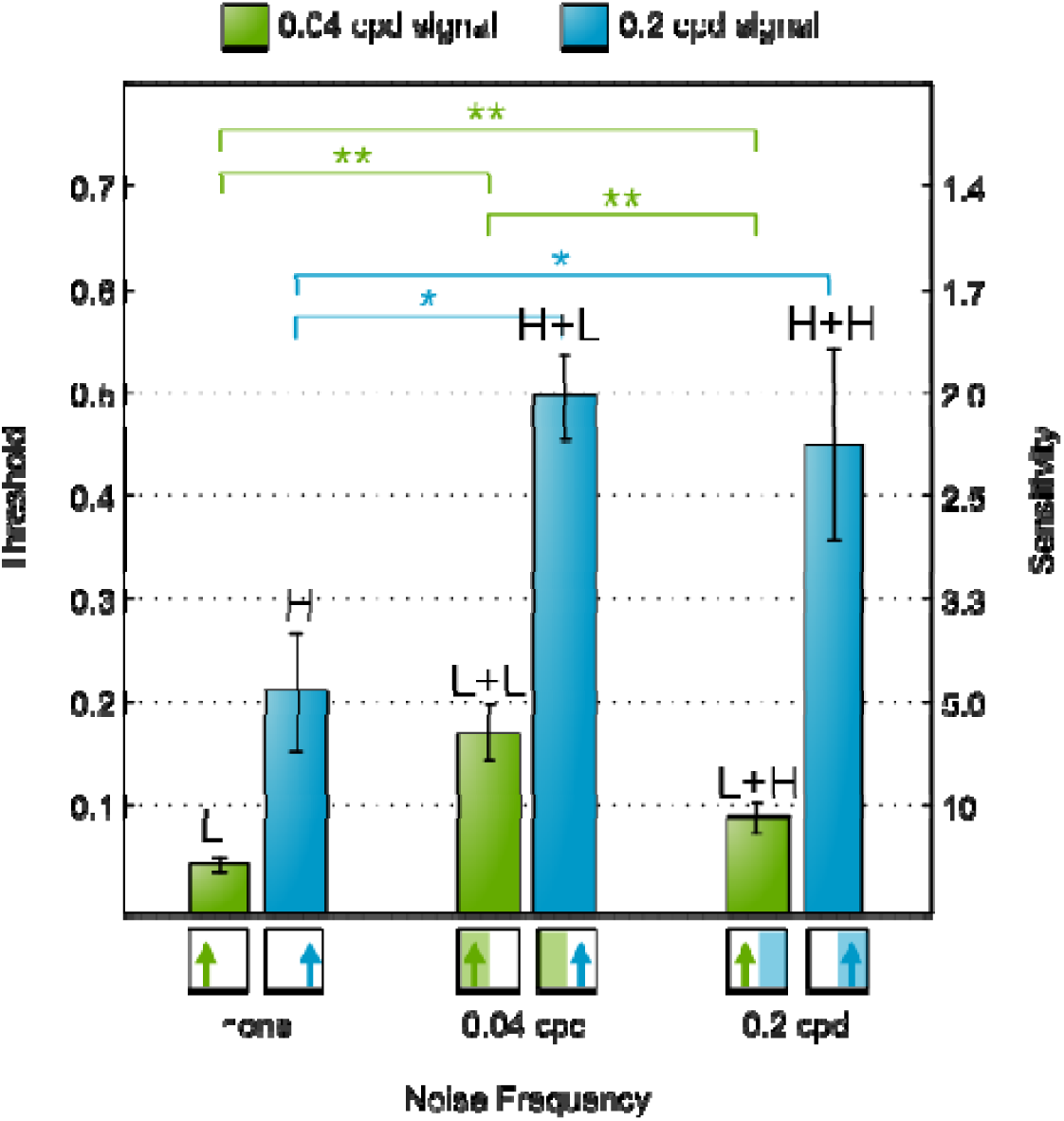
Mantis motion detection contrast thresholds for different combinations of signal and noise frequencies (measured in Experiment M2). Bars show mean contrast detection thresholds for 6 insects and error bars show ± standard error of the mean. Horizontal brackets indicate threshold pairs that differ significantly (paired-sample t-test, * p ≤ 0.05 and ** p ≤ 0.01). Diagrams below bars indicate stimuli conditions as in Fig 2. The results show the same qualitative differences observed in Experiment M1 (Fig 4): the 0.2 cpd signal is masked to similar degrees by noise at either frequency while the 0.04 cpd signal is masked more strongly by 0.04 cpd noise. This similarity excludes the possibility that mantis and human results were different because stimuli appeared spatially distorted to mantises. Letters above bars indicate signal and noise conditions in the format Signal + Noise (S + N). L indicate low spatial frequency, H indicates high spatial frequency. So for example L + H indicates low frequency signal and high frequency noise.

## References

Anderson SJ, Burr DC (1985) Spatial and temporal selectivity of the human motion detection system. Vis Res 25:1147–1154. doi: 0042-6989(85)90104-X [pii]

Anderson SJ, Burr DC (1987) Receptive field size of human motion detection units. Vis Res 27:621–635. doi:0042-6989(87)90047-2[pii]

Anderson SJ, Burr DC (1989) Receptive field properties of human motion detector units inferred from spatial frequency masking. Vis Res 29:1343–1358.

Bahl A, Ammer G, Schilling T, Borst A (2013) Object tracking in motion-blind flies. Nat Publ Gr 16:730–738. doi:10.1038/nn.3386

Borst A (2014) Fly visual course control: behaviour, algorithms and circuits. Nat Rev Neurosci 15:590–9. doi:10.1038/nrn3799

Borst A, Bahde S (1988) Visual information processing in the fly's landing system. J Comp Physiol A 163:167–173. doi:10.1007/BF00612426

Brainard DH (1997) The Psychophysics Toolbox. Spat Vis 10:433–436.

Hassenstein B, Reichardt W (1956) Systemtheoretische analyse der zeit-, reihenfolgen-und vorzeichenauswertung bei der bewegungsperzeption des rüsselkäfers chlorophanus. Zeitschrift für Naturforsch B 11:513–524.

Kleiner M, Brainard D, Pelli D, et al (2007) What's new in Psychtoolbox-3. Perception 36:1.

Lu Z-L, Dosher B (2014) Visual psychophysics: From laboratory to theory. MIT Press, Cambridge, Massachusetts

Moore, BCJ (2012) An Introduction to the psychology of hearing. Emerald group publishing limited. Bingley, UK.

Nityananda V, Tarawneh G, Jones L, et al (2015) The contrast sensitivity function of the praying mantis Sphodromantis lineola. J Comp Physiol A Neuroethol Sens Neural Behav Physiol 201:741–50. doi:10.1007/s00359-015-1008-5

Nityananda V, Tarawneh G, Errington S, Serrano-Pedraza I, Read J (2017) - The optomotor response of the praying mantis is driven predominantly by the central visual field. J Comp Physiol A 203:77–87.

Pelli DG, Brainard DH (1997) The VideoToolbox software for visual psychophysics: Transforming numbers into movies. Spat Vis 10:433–436.

Reichardt W, Wenking H (1969) Optical detection and fixation of objects by fixed flying flies. Naturwissenschaften 56:424–5.

Riehle A, Franceschini N (1984) Motion detection in flies: parametric control over ON-OFF pathways. Exp brain Res 54:390–394.

Srinivasan M V, Lehrer M, Kirchner WH, Zhang SW (1991) Range perception through apparent image speed in freely flying honeybees. Vis Neurosci 6:519–535.

Tarawneh G, Nityananda V, Rosner R, et al (2017) When invisible noise obscures the signal: the consequences of nonlinearity in motion detection. Scientific Reports, in press (preprint available at http://biorxiv.org/content/early/2017/01/05/098459)

